# NetPolicy-RL: Network-Informed Offline Reinforcement Learning for Pharmacogenomic Drug Prioritization

**DOI:** 10.64898/2026.01.29.702506

**Authors:** Ekarsi Lodh, Shalini Majumder, Tapan Chowdhury, Manashi De

## Abstract

Large-scale pharmacogenomic screens provide extensive measurements of drug response across diverse cancer cell lines; however, most computational approaches emphasize point-wise sensitivity prediction or static ranking, which are poorly aligned with practical decision-making, where only a limited number of candidate drugs can be tested. We propose NetPolicy-RL, a biologically informed and decision-centric framework for pharmacogenomic drug prioritization that integrates network diffusion modeling with offline reinforcement learning. Drug selection for each cell line is formulated as an offline contextual bandit problem, enabling direct optimization of ranking quality rather than surrogate regression objectives. Mechanistic biological context is incorporated by propagating drug targets over curated interaction networks (STRING and Reactome) using random walk with restart, and combining the resulting diffusion profiles with cell-specific molecular importance derived from multi-omics data to compute network disruption scores. These biologically grounded signals are integrated with normalized drug response measurements to construct a joint state representation, which is optimized using an offline actor–critic architecture. Across held-out test splits, NetPolicy-RL consistently outperforms global ranking heuristics and learning-to-rank baselines, achieving statistically significant improvements in per-cell Normalized Discounted Cumulative Gain (NDCG@10) and substantial reductions in per-cell regret. Relative to GlobalTopK, the policy improves NDCG@10 for 88.7% of cell lines, while improvements exceed 95% compared with LambdaMART and regression-to-ranking baselines. Ablation analyses show that neither empirical response signals nor network-derived features alone are sufficient, and that their integration yields the most robust performance. Overall, this study demonstrates that combining mechanistic network biology with offline policy learning provides an effective and interpretable framework for drug prioritization in precision oncology.

## 1 Introduction

Precision oncology seeks to align therapeutic agents with the molecular characteristics of individual tumors. Advances in high-throughput screening and molecular profiling have enabled systematic measurement of drug response across diverse cancer cell lines, generating large paired datasets of genomic context and pharmacological outcomes. These resources have driven the development of computational models for drug response prediction and prioritization, intending to inform personalized treatment strategies.

Early computational approaches primarily relied on feature-based regression, mapping gene expression, mutation status, or copy-number alterations to continuous sensitivity measurements using linear or kernel-based models [1], [2]. More recently, deep learning methods have been proposed to capture nonlinear relationships between molecular profiles and drug response, often through multi-omics integration [3], [4], [5]. In parallel, network-based approaches have demonstrated that projecting drug targets and molecular perturbations onto protein–protein interaction or signaling networks can capture pathway-level effects not apparent from gene-level features alone, improving biological interpretability [6], [7].

Reinforcement learning (RL) offers a principled framework for decision-making under uncertainty and has been widely applied in recommender systems and resource allocation. Contextual bandit formulations are particularly well suited to pharmacogenomic settings, as each cell–drug interaction corresponds to a single decision with an immediate outcome. Although still underexplored in biomedical applications, recent studies suggest the potential of contextual bandits for treatment recommendation and adaptive experimental design [8], [9].

A central challenge in pharmacogenomics is that data are inherently offline: experimental outcomes are fixed, costly to obtain, and cannot be adaptively sampled. This precludes standard online RL approaches and necessitates offline policy learning methods that remain stable under distributional shift and limited action coverage, while maintaining alignment between model objectives, biological representations, and evaluation metrics.

In this work, we propose NetPolicy-RL, a hybrid framework that integrates biological network diffusion with offline actor–critic policy learning to directly optimize drug ranking for individual cancer cell lines. Rather than predicting absolute sensitivity values, NetPolicy-RL learns a stochastic drug-selection policy that minimizes regret and maximizes top-*k* ranking quality by combining (i) empirical drug response measurements, (ii) cell-specific molecular context from multi-omics data, and (iii) mechanistic network disruption scores derived from propagating drug targets over curated interaction networks. By embedding network-derived signals into a contextual bandit formulation, the proposed approach aligns optimization with clinically relevant ranking objectives such as NDCG and top-*k* regret, while enabling stable offline learning through an actor–critic architecture with biologically grounded intermediate representations.

The contributions of the paper can be summarized as follows:

i. Formulation of drug prioritization in pharmacogenomics as an offline contextual bandit problem focused on ranking rather than prediction.
ii. Introduction of a principled integration of network diffusion-based disruption modelling into policy learning.
iii. Demonstration of consistent improvements over static and learning-to-rank baselines across multiple evaluation metrics.
iv. Providing mechanistic insights into policy behavior through oracle-aligned ranking analyses.

The rest of the paper follows the following arrangement: Section 2 summarizes the recent advances in the research in the related field, as in our study. Section 3 encompasses those fundamental concepts that are covered in our study. Methods and materials required for the study fall under Section 4. The result analysis portion is shown in Section 5. The conclusion of our study is drawn in Section 6.

## 2 Related Studies

A substantial body of work models cancer drug response by learning joint representations of cell lines and compounds, typically optimizing pointwise regression losses (e.g., AUC/IC50) [10], [11] or pairwise ranking surrogates. Early deep models such as CDRscan demonstrated that deep neural networks can map genomic features to drug effectiveness at scale [12]. More recent methods increasingly use structure-aware drug encoders, for example graph neural networks that represent compounds as molecular graphs and fuse them with cell features. DeepCDR integrates multi-omics cell profiles with graph-based drug representations to predict response, highlighting the value of combining multiple molecular modalities [13]. GraphDRP similarly models drugs as graphs and cell lines as genomic features, showing strong performance in response prediction [14].

Beyond pure prediction, interpretability and representation learning have become important. The PaccMann line of work explores multimodal attention mechanisms for anticancer sensitivity prediction and provides a web service for model access and interpretability [3], [15]. Graph-based and contrastive learning approaches such as GraphCDR aim to improve generalization by adding auxiliary representation objectives [16]. Complementary developments include graph-structured cell encoders that incorporate PPI networks into the modeling pipeline (e.g., DrugGCN) to capture pathway or sub-network effects [17]. Taken together, many methods optimize prediction losses rather than decision-oriented top-*k* prioritization.

Network propagation and diffusion methods are widely used for drug repositioning, and target identification, motivated by the observation that disease modules and drug effects manifest at pathway/network scales rather than single genes. Recent studies have also highlighted network-based reasoning (including diffusion/random-walk variants) as a core family of computational repurposing strategies [18], [19], [20]. Empirically, network propagation of genetic evidence has been shown to support drug target identification, demonstrating the practical utility of diffusion-style scoring [21]. More recent multilayer extensions (e.g., random-walk-with-restart variants on multilayer networks) further illustrate how diffusion can prioritize genes/drugs by integrating heterogeneous biological relations [22]. In addition, modern target-prioritization ecosystems such as the Open Targets Platform reflect the field-wide emphasis on evidence integration for target discovery and prioritization [23].

The feasibility of large-scale computational drug prioritization depends on expansive screening resources. The PRISM Repurposing dataset provides viability profiling for thousands of compounds across hundreds of cancer cell lines and is publicly available through the DepMap portal, making it a key benchmark source for repurposing-style prioritization [24]. Parallel efforts to build and harmonize dependency maps and multi-omic annotations further strengthen the ecosystem for systematic evaluation and mechanistic interpretation [25], [26].

A complementary research direction treats biomedical recommendation and treatment selection as policy learning from logged data. Offline RL and contextual bandits are especially relevant when online exploration is infeasible or unsafe. Practical challenges such as policy evaluation and model selection are well documented in healthcare-focused offline RL analyses [27]. Broader surveys also summarize how RL is used across healthcare decision problems and highlight the importance of evaluation rigor under offline assumptions [28], [29], [30]. Recent work on off-policy optimization in contextual bandits and offline settings provides algorithmic foundations for learning policies from fixed datasets [31], [32].

Contextual bandits represent a simplified, single-step variant of reinforcement learning and have been widely applied to recommendation and treatment selection problems. In healthcare and related domains, offline contextual bandits have been used to model decision-making under uncertainty, where the objective is to optimize top-*k* utility rather than predictive accuracy [27], [33]. Recent work has further explored conservative and variance-aware learning principles to improve robustness in offline settings [34], [35].

While offline RL and contextual bandits have been extensively studied in recommender systems and healthcare decision support, their application to pharmacogenomic drug prioritization remains limited. In particular, existing drug response models rarely adopt decision-centric formulations that directly optimize ranking quality. The present work builds on these foundations by integrating offline policy learning with biologically informed network modeling to address drug prioritization under realistic experimental constraints.

## 3 Preliminaries

### 3.1 Drug Ranking in Pharmacogenomics

In pharmacogenomic screens, each cancer cell line is tested against multiple drugs, producing a set of observed responses. For a given cancer cell line *c*, let

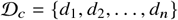

denote the set of drugs tested on that cell line, and let *r* (*c, d)* denote the observed or derived reward associated with drug *d*. A drug recommendation model induces a ranking over 𝒟_*c*_, and evaluation assesses how well this ranking aligns with the empirical response landscape, particularly at the top of the list.

### 3.2 Oracle Ranking and Oracle Rank [33]

For each cell line *c*, the oracle ranking is defined by sorting drugs in descending order of their observed rewards *r* (*c, d*). The oracle rank of a recommended drug *d* is given by:

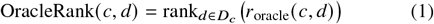

where smaller values correspond to better-performing drugs. Oracle rank provides a direct reference for evaluating how close a model’s recommendations are to the optimal choice.

### 3.3 Normalized Discounted Cumulative Gain (NDCG) [36]

Normalized Discounted Cumulative Gain (NDCG) evaluates how well effective drugs are ranked near the top of a recommendation list, while accounting for the position of each drug. The discounted cumulative gain at rank *k* for cell line *c* is defined as:

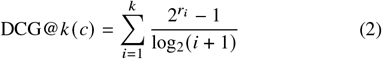

where *r*_*i*_ denotes the reward of the drug ranked at position *i*. The normalized score is given by:

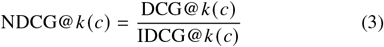

where IDCG@*k* (*c)* denotes the DCG of the oracle ranking. NDCG values lie in [0, 1], with higher values indicating better ranking quality.

### 3.4 Top-*k* Regret [30]

Top-*k* regret (regret@k) measures the performance loss incurred when the best possible drug is not included among the top-*k* recommendations. The top-*k* regret for a cell line *c* is defined as:

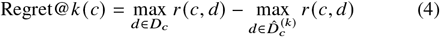

where 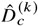 denotes the set of top-*k* drugs recommended by the model. Lower regret indicates that near-optimal drugs are included among the top recommendations.

### 3.5 Hit Rate at *k* [37]

Hit@*k* evaluates whether at least one highly effective drug appears among the top-*k* recommended candidates for a given cell line.

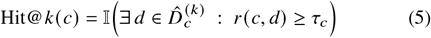

where *τ*_*c*_ denotes a cell-specific relevance threshold, such as a top quantile of observed rewards.

## 4 Methods and Materials

### 4.1 Methodological Overview

This work proposes NetPolicy-RL, a hybrid framework that integrates biological network diffusion with offline reinforcement learning to prioritize anticancer drugs for a given cellular context. Instead of focusing on predicting scalar sensitivity values (e.g., IC50) for individual cell-drug pairs, our objective is to learn a drug-selection policy that directly ranks candidate drugs for each cell line.

The framework consists of two tightly coupled layers:

1. **Biology-informed Network diffusion layer:** Models how drug targets propagate influence across a protein-protein interaction (PPI) or signaling network to compute cell-specific network disruption score.
2. **Decision-oriented Policy learning layer:** Formulated as a contextual bandit problem in a reinforcement learning setup, which learns a stochastic policy over drugs that maximizes expected therapeutic reward using offline pharmacogenomic data.

The overall methodology is depicted in Fig. 1.

**Fig. 1:**
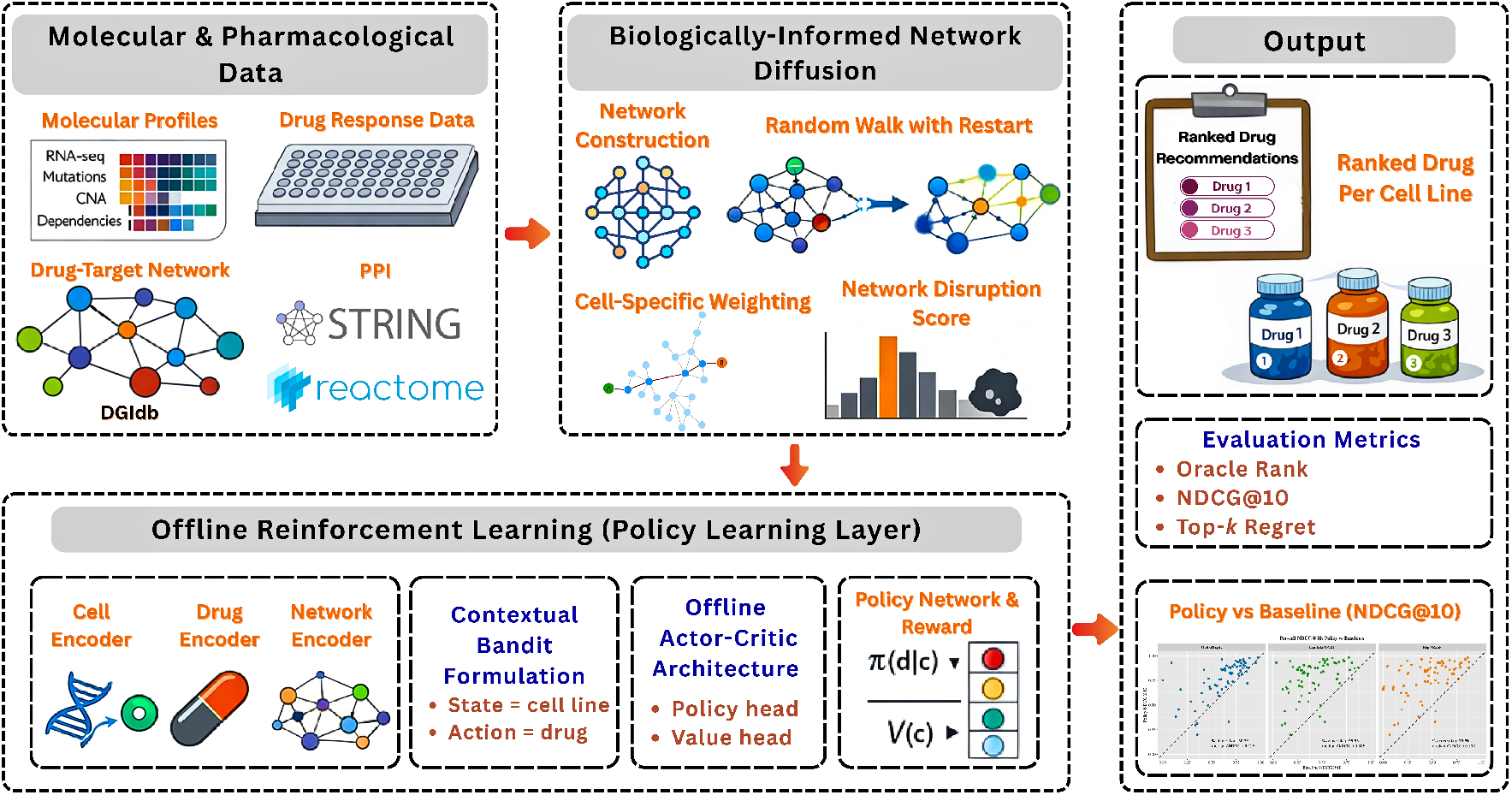
Overall methodology of NetPolicy-RL for pharmacogenomic drug prioritization. The workflow integrates molecular, pharmacological, and network-level information with offline reinforcement learning to derive cell-specific drug rankings. (Left) Multi-omics molecular profiles of cancer cell lines (RNA-seq expression, mutations, copy-number alterations, and gene dependencies) together with drug response measurements are collected, and drug–target associations are obtained from DGIdb. These are mapped onto curated biological interaction networks from STRING and Reactome. (Center) A biologically informed network diffusion layer propagates drug target influence over the interaction network using random walk with restart, combined with cell-specific gene importance weighting to compute network disruption scores for each cell–drug pair. (Bottom) An offline reinforcement learning layer formulates drug selection as a contextual bandit problem, where the state corresponds to a cell line and the action to a drug. Cell, drug, and network encoders produce a joint representation that is optimized using an offline actor–critic architecture with policy and value heads, guided by a reward integrating normalized drug response and network disruption. (Right) The learned policy outputs ranked drug recommendations for each cell line, which are evaluated using decision-centric metrics, including oracle rank, NDCG@10, and top-k regret, and compared against baseline ranking methods.

### 4.2 Molecular and Pharmacological Data Sources

#### 4.2.1 Cancer Cell Line Molecular Profiles

Cell line molecular data are obtained from the DepMap portal [38], [39], [40], which provides harmonized multi-omics measurements across hundreds of cancer cell lines. For each cancer cell line, the following gene-level modalities are used:

- Gene expression (RNA-Seq)
- Somatic mutation status
- Copy number alterations (CNA)
- CRISPR gene dependency scores

For a given cell line *c*, molecular information is represented at the gene level. Let *V* and *F* denote the set of genes present in the interaction network and the number of molecular features, respectively. Each cell line is associated with a feature matrix:

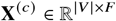

where each row corresponds to a gene, and each column corresponds to a molecular feature. This representation preserves gene-level resolution and enables direct integration with network-based modeling.

#### 4.2.2 Drug Response Measurements

Drug sensitivity data are collected from large-scale pharmacogenomic screens, including GDSC [1], [41], [42] and PRISM Repurposing. These datasets report quantitative measures of drug efficacy, such as IC50 or area under the dose-response curve (AUC), across many cell-drug combinations.

To ensure comparability across drugs and datasets, raw response values are normalized on a per-drug basis:

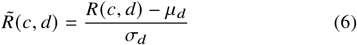

where *μ*_*d*_ and *σ*_*d*_ denote the mean and standard deviation of responses for drug *d*. This prevents drugs with inherently wider response ranges from dominating the learning objective.

#### 4.2.3 Drug-Target Associations

Each drug is mapped to its known molecular targets using curated drug-gene interactions from DGIdb [43]. For a drug *d*, the target set is defined as:

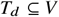

where *V* is the gene set of the interaction network. These target sets serve as the biological entry points for network diffusion and encode prior mechanistic knowledge about the action of drugs.

### 4.3 Network Diffusion-Based Disruption Modeling

#### 4.3.1 Interaction Network Construction

A global molecular interaction network is constructed to model intracellular signal propagation. The network is represented as a graph *G* = (*V, E*), where nodes correspond to genes and edges represent protein-protein or signaling interactions.

Two complementary network backbones are considered: the STRING protein-protein interaction network [44] and curated signaling pathways from Reactome [45], [46]. These networks provide complementary views of molecular connectivity, with STRING emphasizing physical interactions and Reactome emphasizing functional signaling relationships. Using multiple backbones allows assessment of robustness across different biological interaction assumptions.

#### 4.3.2 Random Walk with Restart (RWR) for Target Propagation

The way in which a drug’s molecular targets influence the broader cellular network, Random Walk with Restart (RWR) is employed. For a given drug *d*, an initial probability vector **p**_0_ is defined over network nodes such that probability mass is uniformly distributed across the drug’s target genes:

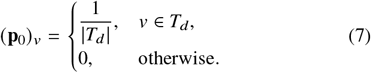

The diffusion process evolves and iteratively updates as follows:

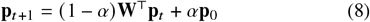

where **W** is the column-normalized adjacency matrix of *G* and *α* ∈ (0, 1) is the restart probability. Iteration continues until convergence, yielding a stationary distribution **p**_∞_. This distribution captures how drug target influence propagates through the molecular network beyond direct targets.

#### 4.3.3 Cell-Specific Node Importance Weighting

While diffusion captures drug-centric influence, cellular context is introduced through gene-specific importance weights. For each gene *v* in the cell line *c*, a weight 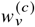 is computed as a deterministic function of molecular measurements:

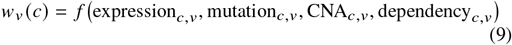

where *f* (·) denotes the deterministic aggregation function.

This weighting reflects the intuition that perturbing a highly expressed or highly essential gene is likely to have greater phenotypic consequences than perturbing a marginally active gene. Importantly, these weights are cell-specific, allowing the same drug to induce different network effects across cell lines.

#### 4.3.4 Network Disruption Score

The final network-level impact of drug *d* on cell line *c* is quantified by the network disruption score:

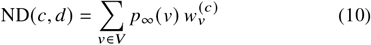

where *p*_∞_ (*v)* denotes how strongly drug targets influence gene *v* and 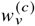 denotes how important gene *v* is in cell line *c*.

This score integrates drug-centric diffusion with cell-centric molecular importance, yielding a scalar measure of how strongly a drug is expected to disrupt critical network regions in a given cellular context. Optionally, disruption scores can be aggregated at the pathway level by summing diffusion mass over predefined pathway gene sets.

This complete pipeline integrating network construction, molecular feature assembly, and diffusion-based network disruption computation is detailed in Algorithm 1.

##### Algorithm 1

Graph and Feature Preprocessing

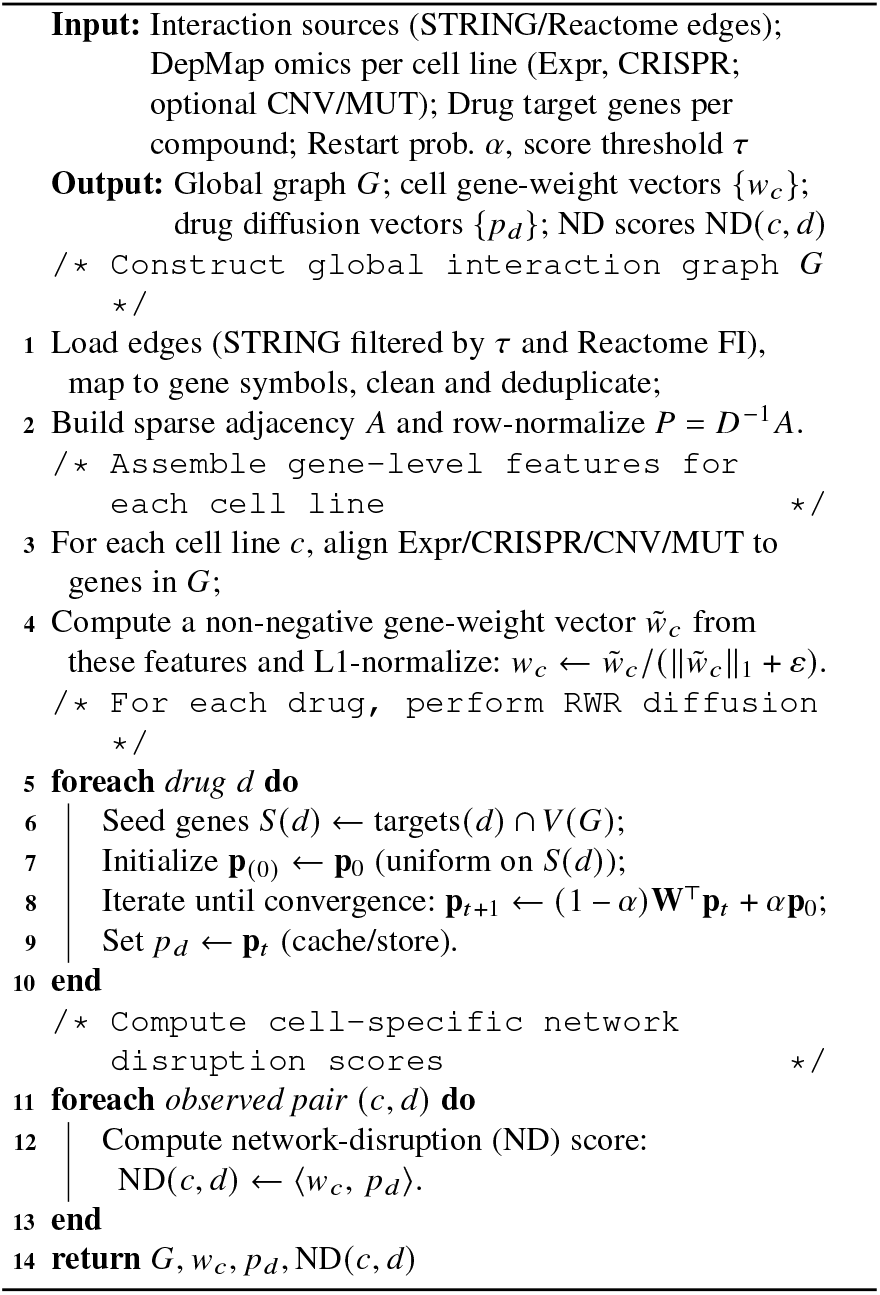

### 4.4 Reinforcement Learning Formulation

#### 4.4.1 Contextual Bandit Perspective

Drug selection is formulated as a contextual bandit problem.

- Each cell line corresponds to a **state** *s*_*c*_.
- Each drug corresponds to an **action** *a*_*d*_.
- Outcome of selecting a drug is summarized by a scalar **reward** *r* (*c, d*).

Unlike multi-step reinforcement learning, this formulation is appropriate because drug response screens provide only single-step outcomes.

#### 4.4.2 Reward Definition

The reward function integrates empirical drug response with mechanistic network information:

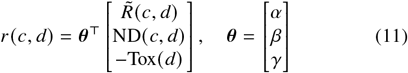

where 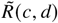, ND(*c, d*) and Tox(*d*) represents the normalized drug sensitivity, network disruption and toxicity penalty respectively, and *α, β* and *γ* are the weighting coefficients controlling the relative contributions of measured sensitivity, network disruption, and toxicity penalty and *α, β, γ* ≥ 0. This formulation encourages the policy to prioritize drugs that are both empirically effective and mechanistically disruptive in relevant cellular networks.

Based on the defined reward formulation, the construction of the offline training dataset used for policy learning is summarized in Algorithm 2.

##### Algorithm 2

Training Sample Generation

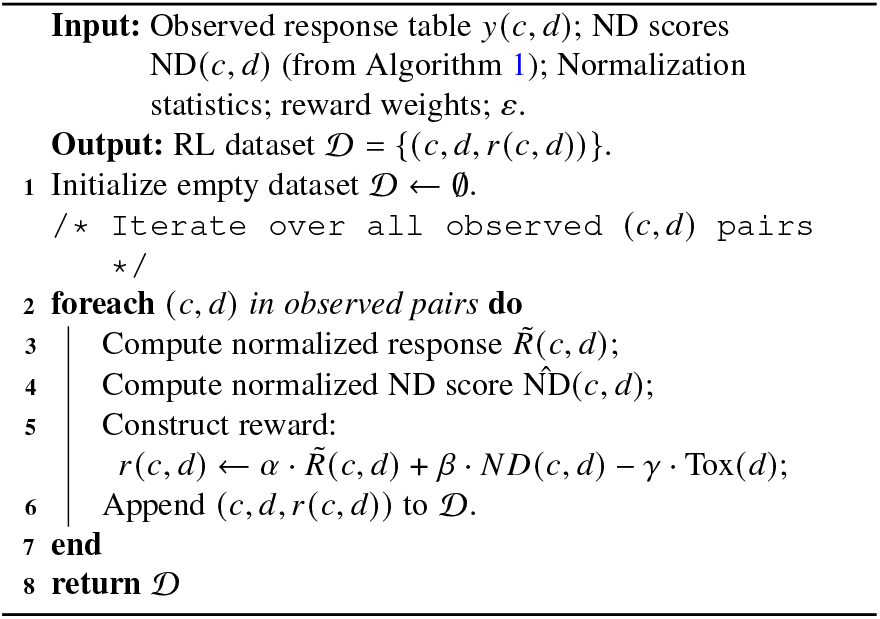

#### 4.4.3 Offline Learning Assumption

The framework operates under an offline reinforcement learning setting, where policy optimization is performed exclusively using previously collected pharmacogenomic data. The agent does not interact with the biological system during training and does not generate new cell-drug experiments.

Formally, learning is based on a fixed dataset:

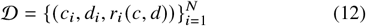

where each tuple corresponds to an observed response of drug *d*_*i*_ on cell line *c*_*i*_. The dataset is static and fully specified prior to training.

Since each decision corresponds to a single drug selection for a given cellular context, the problem reduces to a contextual bandit formulation with no state transitions. Consequently, the return for each decision is equal to the immediate reward, and no temporal discounting is required.

This offline formulation reflects practical experimental constraints in pharmacogenomics and ensures that the learned policy is grounded entirely in observed biological evidence.

### 4.5 Model Architecture

#### 4.5.1 Cell Encoder

Cell encoder maps the deterministically computed molecular state of a cancer cell line into a compact latent representation that captures both gene-level activity and network context. For a given cell line *c*_*i*_, the encoder takes as input the interaction network *G* = (*V, E)* and the cell-specific molecular feature matrix **X**^(*c*)^ as defined in Section 4.2.1.

Gene-level molecular measurements are aggregated into a cell-specific importance vector *w* ^(*c*)^. This vector reflects the relative functional importance of genes within the cellular context. To obtain a fixed-dimensional representation suitable for downstream learning, the cell-specific gene importance vector *w* ^(*c*)^ is projected onto the top *D*_*c*_ principal components learned from the training cell lines. The PCA is subsequently applied to all cell lines to prevent information leakage. The final cell embedding is given by:

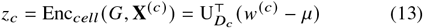

where *μ* denote the mean gene-importance vector and 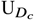 the matrix of leading principal directions. This embedding encodes pathway-level molecular vulnerabilities while preserving biological network structure.

#### 4.5.2 Drug Encoder

The drug encoder produces a compact representation of each compound that captures its pharmacological properties. Each drug *d* is represented by a fixed-length descriptor vector derived from available drug metadata.

The encoder applies a deterministic feature mapping to obtain a drug embedding:

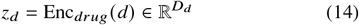

This formulation allows the policy to generalize across compounds and supports evaluation on drugs not explicitly observed during training, provided corresponding descriptors are available.

#### 4.5.3 Network Impact Encoder

The network impact encoder summarizes the effect of a drug on the molecular network of a given cell line. It is implemented as an identity mapping:

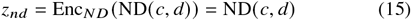

This scalar embedding offers a biologically interpretable measure of network-level drug impact, complementing molecular and pharmacological features in the state representation.

#### 4.5.4 Policy and Value Heads

The output of the cell, drug, and network impact encoders are concatenated to form a joint representation for each cell-drug pair:

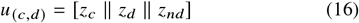

This combined representation captures complementary aspects of the decision problem: the cellular context, the drug’s intrinsic properties, and the expected biological impact of the drug on the cell. Two neural heads process this shared representation within an actor-critic framework. The policy head, parameterized by ***ψ***, produces a probability distribution over candidate drugs for a given cell line:

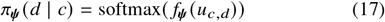

This distribution defines the drug-selection policy and directly determines the ranking of candidate therapies.

In parallel, the value head, parameterized by ***ϕ***, estimates the expected reward associated with the cellular context:

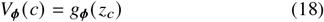

The value function provides a baseline estimate of achievable benefit and is used to stabilize policy optimization. The separation of policy and value estimation is a defining characteristic of actor-critic methods and is essential for reducing the overall variance during learning.

### 4.6 Offline Actor-Critic Optimization

Policy learning is performed using an offline actor-critic optimization strategy. The objective is to learn a stochastic drug-selection policy parameter ***ψ*** that maximizes expected reward using a fixed dataset of previously observed cell-drug responses, reflecting the practical constraints of drug response modeling, where experimental outcomes are fixed and costly to obtain.

Each observed cell-drug pair corresponds to a single decision step, consistent with a contextual bandit setting. The optimization objective is to learn policy parameters ***ψ*** that maximize expected reward under the empirical data distribution, while simultaneously learning a value function *V*_***ϕ***_ that approximates expected outcomes for each cell line.

#### 4.6.1 Advantage Estimation

To access the relative quality of selecting a specific drug *d* for a given cell line *c*, an advantage function is defined as:

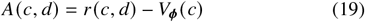

The advantage measures how much better or worse the observed reward for drug *d* is compared to the expected reward baseline for cell *c*. Positive advantages indicate drugs that outperform expectations, while negative advantages indicate suboptimal choices.

This formulation allows the policy to focus on relative improvements rather than absolute reward magnitudes.

#### 4.6.2 Loss Functions

##### Policy Loss

The policy network is optimized using a REINFORCE-style objective with a value baseline by maximizing the expected advantage-weighted log-likelihood of selected actions. The policy loss is defined as:

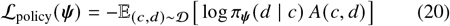

This loss increases the probability of selecting drugs that yield higher-than-expected rewards and decreases the probability of selecting drugs with poor outcomes.

##### Value Loss

The value function is trained to minimize the discrepancy between predicted and observed rewards using a mean-squared error objective:

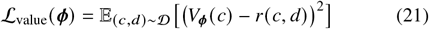

Accurate value estimation is crucial, as errors in the value function directly affect advantage computation and, consequently, policy updates.

##### Total Loss

The overall optimization objective combines both components:

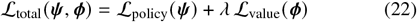

where *λ* controls the relative contribution of value learning. This joint optimization ensures that the policy improves while maintaining a stable and informative baseline.

The overall actor-critic optimization and policy training is summarized in Algorithm 3.

### 4.7 Ablation Study Design

To assess the contribution of network-derived signals, we performed a controlled ablation study in which the learning architecture and optimization procedure are kept fixed, and only the context feature set provided to the policy and value networks is varied. All ablated models are trained and evaluated using identical data splits, normalization procedures, and ranking metrics.

Specifically, we evaluate the following variants:

i. **Response-only:** using summary statistics of observed drug responses per cell line.
ii. **STRING-only:** using summary statistics of diffusion-based network disruption scores computed on the STRING interaction network.
iii. **Reactome-only:** using summary statistics of diffusion-based network disruption scores computed on the Reactome network; and
iv. **Full:** combining response-based and network-based features.

#### Algorithm 3

Offline Actor-Critic Training Loop

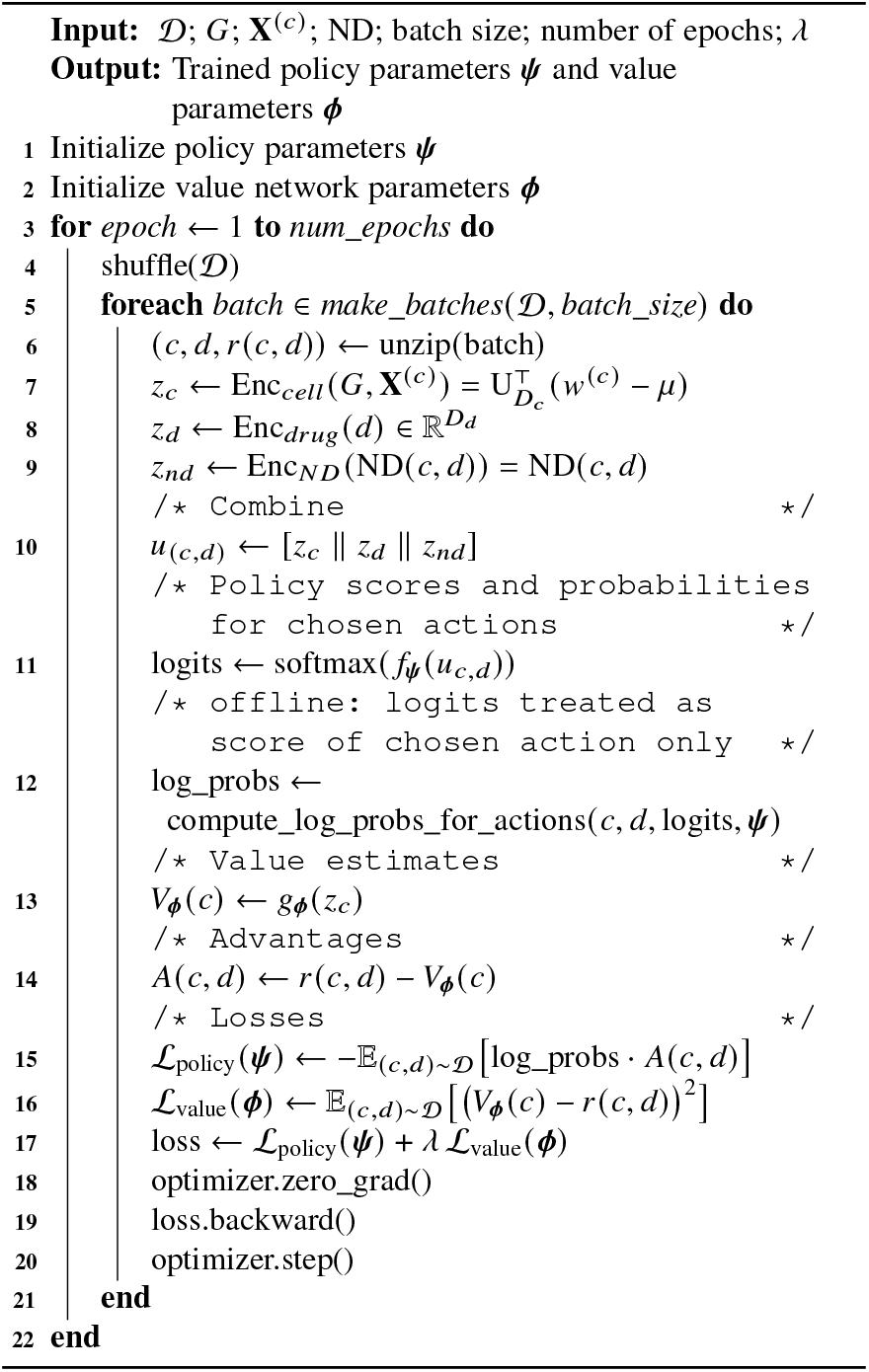

Performance comparisons across variants isolate the impact of network diffusion and interaction backbone choice on policy learning.

## 5 Result Analysis

### 5.1 Per-Cell Regret Analysis

We first assess the effectiveness of the learned policy in reducing per-cell regret, which measures the gap between the best achievable drug response and the best response among the top-*k* recommended drugs for each cell line. Fig. 2 shows the distribution of per-cell regret for the baseline ranking strategy and the learned policy.

**Fig. 2:**
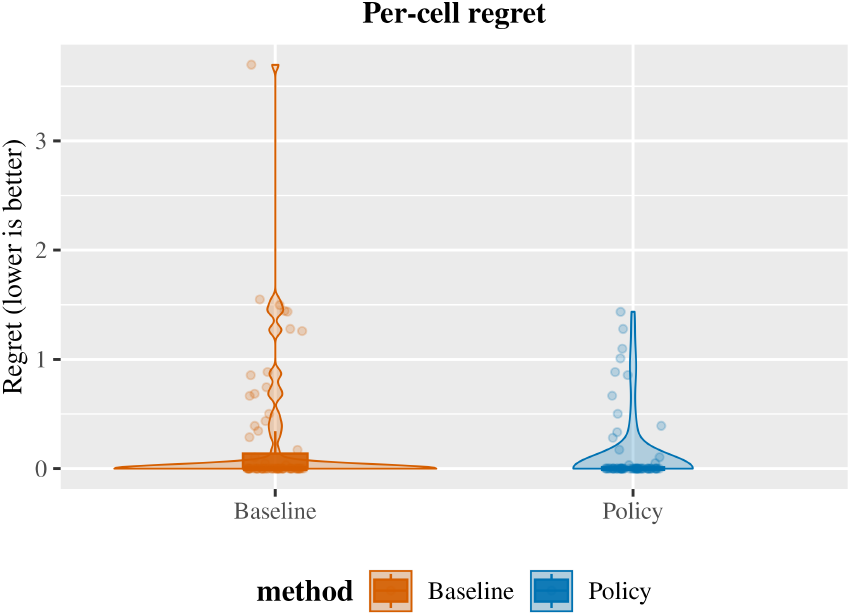
Distribution of per-cell regret for the baseline and policy-based ranking methods. Lower values indicate better performance

The baseline exhibits a broad and heavy-tailed distribution, with several cell lines incurring large regret values. This indicates that the baseline frequently fails to include near-optimal drugs among its top recommendations, particularly for more challenging cellular contexts.

In contrast, the policy produces a markedly tighter regret distribution concentrated near zero. Both the median regret and the upper tail are substantially reduced, demonstrating that the policy consistently selects drugs closer to the oracle-optimal choice across cell lines. Importantly, the reduction is not limited to a small subset of cases: the overall compression of the distribution indicates improved robustness rather than isolated gains.

The suppression of extreme regret values is particularly notable, as it suggests that the policy avoids catastrophic misranking for difficult cell lines. This behavior is critical in pharmacogenomic decision-making, where poor recommendations for a subset of cases can dominate overall performance.

These results demonstrate that framing drug prioritization as a policy learning problem leads to systematic improvements in decision quality compared with static baseline ranking approaches.

### 5.2 Paired Cell-Line Comparison

Paired comparisons were performed to assess whether improvements in ranking quality occur consistently at the level of individual cell lines. For each cell line, performance under the baseline and the learned policy was compared using both per-cell regret and per-cell NDCG@10.

Fig. 3 presents paired results for the two metrics. In panel (a), per-cell regret values under the baseline and policy are connected for each cell line. The majority of trajectories exhibit a clear downward shift, indicating reduced regret under the policy. A Wilcoxon signed-rank test confirms that this reduction is statistically significant (*p* = 8.30 ×10^*−*3^), demonstrating that the policy consistently selects drugs closer to the oracle-optimal choice across cell lines.

**Fig. 3:**
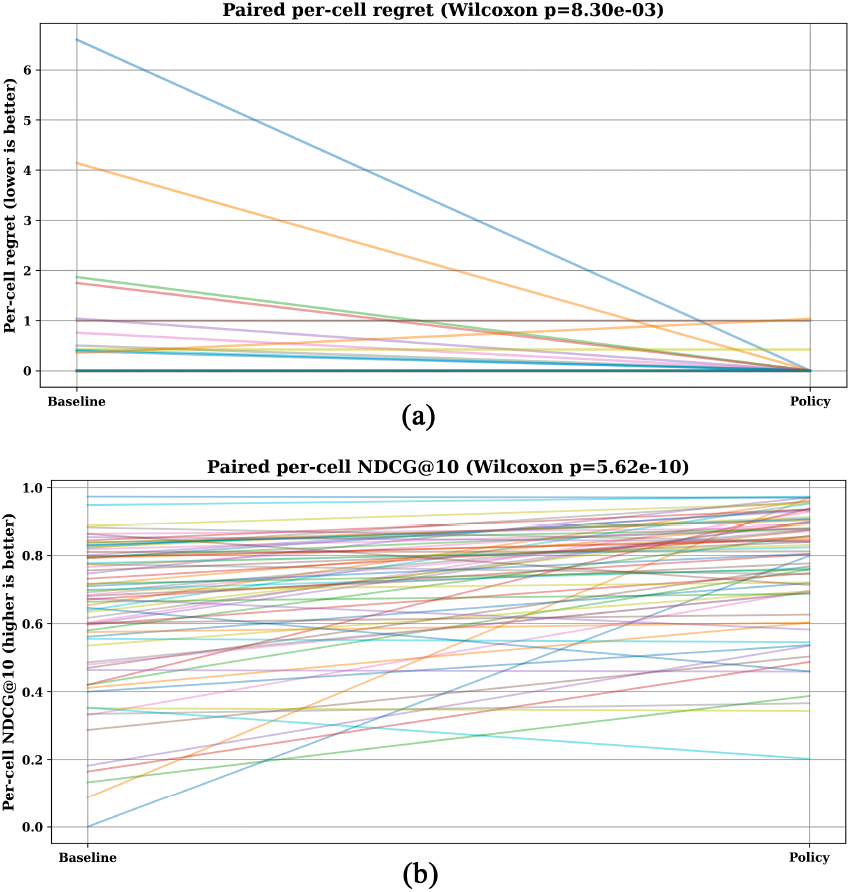
Paired per-cell comparison between baseline and policy. (a) Per-cell regret (lower is better). (b) Per-cell NDCG@10 (higher is better). *p*-values are from two-sided Wilcoxon signed-rank tests

In panel (b), paired per-cell NDCG@10 values are shown. Most trajectories slope upward, reflecting improved ranking quality under the policy. The increase in NDCG@10 is highly significant according to the Wilcoxon signed-rank test (*p* = 5.62 × 10^*−*10^), indicating that effective drugs are systematically promoted toward the top of the ranked lists.

Taken together, the paired analyses demonstrate that the gains achieved by the policy are not driven by aggregate effects or a small subset of favorable cases. Instead, improvements in both regret and ranking quality are observed consistently across individual cell lines, supporting the robustness of the learned decision policy.

### 5.3 Training Dynamics and Optimization Stability

To examine optimization behavior, we analyze the training and validation loss curves obtained during offline actor-critic learning.

Fig. 4 shows the evolution of training and validation losses across epochs. Both curves decrease sharply during early training, followed by gradual convergence to stable plateaus. Importantly, the validation loss closely tracks the training loss throughout optimization, with no evidence of divergence or instability.

**Fig. 4:**
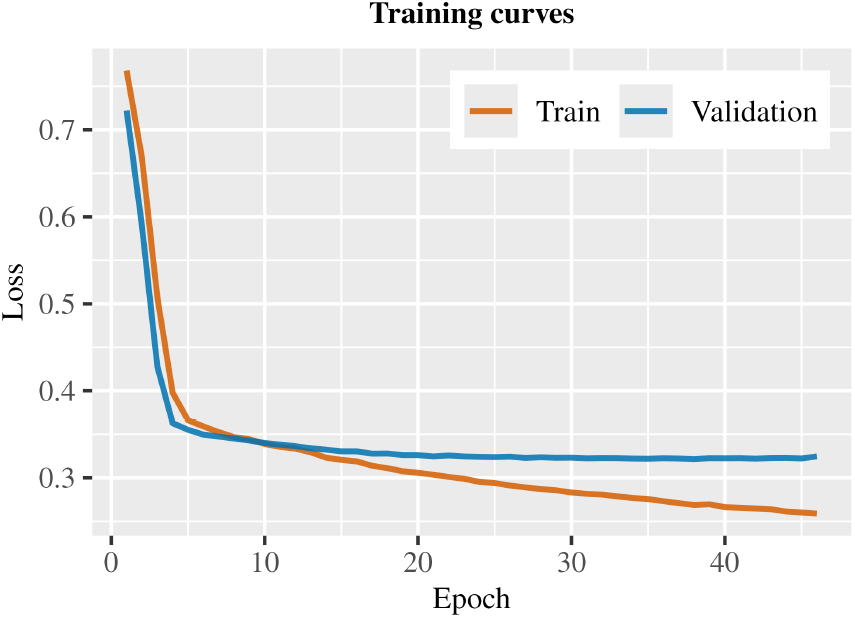
Training and validation loss curves for offline actor–critic optimization

This behavior highlights two key aspects of the proposed framework. First, the use of a value function baseline (Section 4.5.4) stabilizes policy updates by reducing gradient variance. Secondly, the offline learning assumption (Section 4.4.3) does not lead to overfitting, despite the fixed dataset, indicating that the learned policy captures generalizable structure across cell lines rather than memorizing observed responses.

### 5.4 Contribution of Network-Derived Features

The contribution of the different components of the state representation were evaluated through a systematic ablation study, as described in Section 4.7.

Fig. 5 summarizes test-set performance across ablation variants using NDCG@*k* and mean regret. The response-only variant achieves moderate ranking performance, indicating that empirical drug sensitivity signals alone provide useful but incomplete guidance for drug prioritization. Both network-only variants improve upon the response-only model in terms of NDCG@*k*, demonstrating that diffusion-based propagation over molecular interaction networks captures biologically meaningful structure relevant to drug response. Between the two, STRING-only and Reactome-only models exhibit comparable performance, suggesting that the observed gains are not specific to a single network source.

**Fig. 5:**
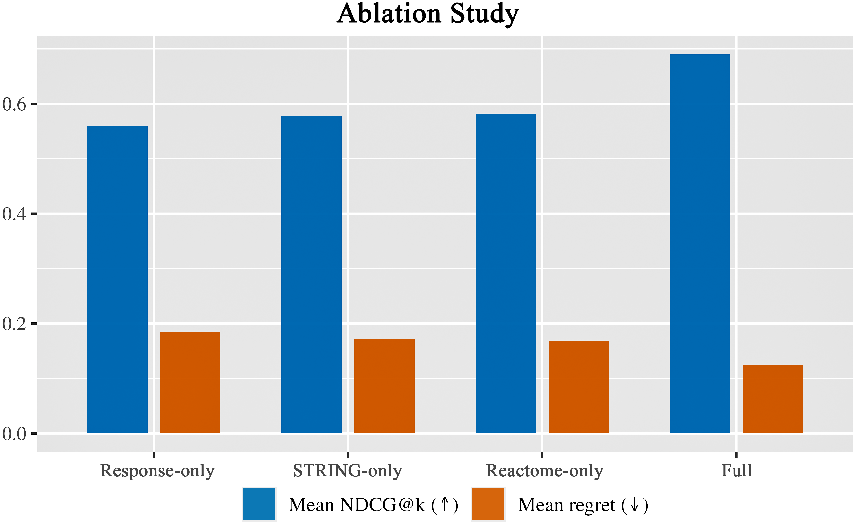
Ablation study comparing response-only, STRING-only, Response-only, and full models

The full model, which integrates normalized response signals with network diffusion features, achieves the highest NDCG@*k* and the lowest mean regret. The reduction in regret is particularly pronounced, indicating that the integrated model more reliably selects drugs closer to the oracle-optimal choice across cell lines. This improvement over both response-only and network-only variants confirms that network-derived information provides complementary signal rather than acting as a surrogate for observed response.

Overall, the ablation results demonstrate that optimal performance arises from the joint use of empirical drug response data and mechanistic network modeling, and that integrating multiple network backbones further stabilizes ranking quality.

### 5.5 Oracle-Based Ranking Behavior

To gain quantitative insights into how the learned policy alters drug rankings, we analyze recommendations relative to oracle rankings, which represent ideal drug orderings based on observed rewards. Fig. 6 shows the oracle top-20 drug rankings for two such cell lines, with points indicating whether a drug is included in the policy’s top-*k* recommendations.

**Fig. 6:**
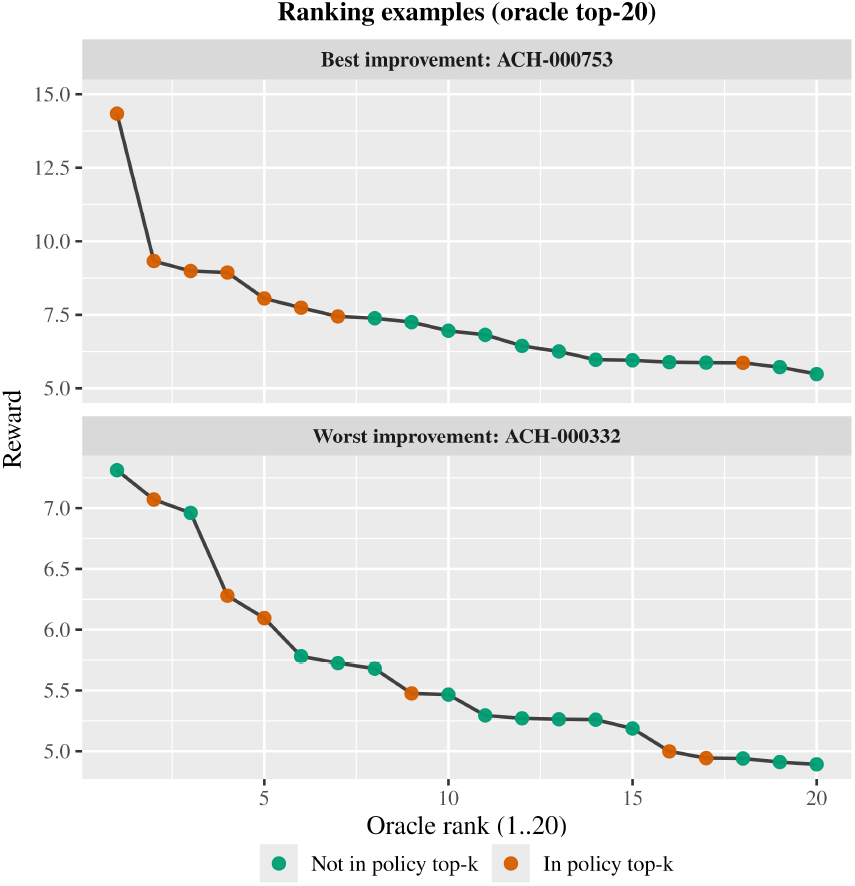
Oracle-aligned ranking examples for representative cell lines. Top: best improvement (ACH-000753). Bottom: worst improvement (ACH-000332). Points indicate whether a drug is included in the policy’s top-*k*recommendations

In the best-improvement case (ACH-000753), drugs selected by the policy are concentrated toward the top of the oracle ranking. Several high-reward compounds within the oracle top-10 are included in the policy’s recommendation set, resulting in a reward profile that closely follows the oracle curve in the high-impact region. This alignment explains the substantial reduction in regret observed for this cell line and illustrates the policy’s ability to prioritize strongly effective drugs when the response landscape is well-structured.

In contrast, the worst-improvement case (ACH-000332) exhibits a flatter reward gradient across the oracle ranking, indicating a more ambiguous response landscape. Although the policy still includes some oracle-top drugs within its top-*k* recommendations, coverage is less concentrated at the very top of the ranking. As a result, the achievable improvement is smaller, despite the policy avoiding consistently low-reward selections. This behavior highlights an inherent limitation imposed by the underlying response distribution rather than a failure of the policy itself.

Overall, these examples demonstrate that the learned policy adapts its ranking behavior to the structure of the oracle reward landscape. When strong signal separation exists, the policy effectively recovers high-reward drugs; when the landscape is less sharply defined, improvements are naturally constrained. This observation is consistent with the quantitative regret and NDCG analyses and underscores the robustness of the policy across heterogeneous cell-line-specific response profiles.

### 5.6 Comparison with Baseline Ranking Methods

The performance of the learned policy was compared against multiple baseline ranking approaches using per-cell NDCG@10, which evaluates the quality of the top-ranked drug recommendations for each cell line.

Fig. 7 presents scatter plots comparing policy NDCG@10 against three baselines: GlobalTopK, LambdaMART [47], and regression-to-ranking (Reg → Rank). Each point corresponds to a cell line, and the dashed diagonal indicates equal performance between the policy and the baseline. Points above the diagonal represent cell lines for which the policy achieves higher NDCG@10. Across all baselines, the majority of points lie above the diagonal, indicating that the policy improves ranking quality for most cell lines. Relative to the GlobalTopK baseline, 88.7% of cell lines show improved NDCG@10 under the policy, with a median increase of + 0.125. This demonstrates that the policy substantially outperforms static global ranking strategies that ignore cell-specific context.

**Fig. 7:**
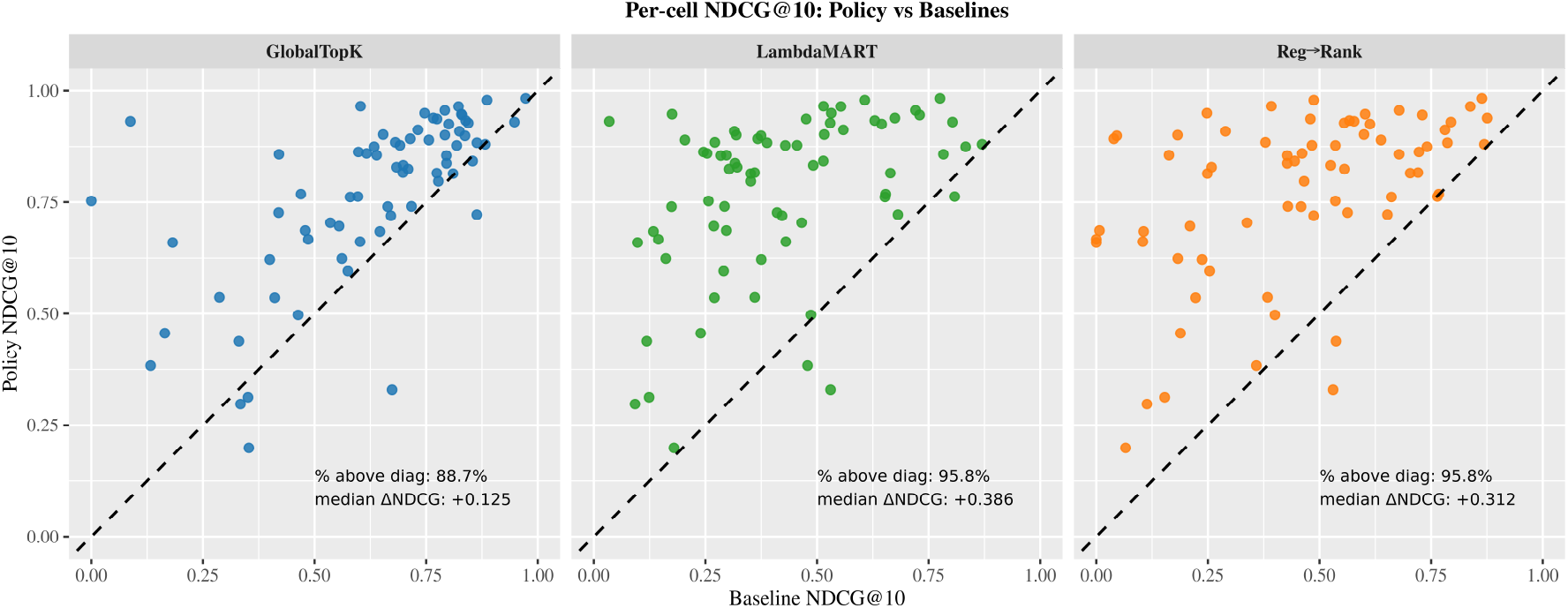
Per-cell NDCG@10 comparison between the learned policy and baseline ranking methods. Points above the diagonal indicate improved ranking under the policy. Annotated percentages and median ΔNDCG values summarize per-cell improvements

The strongest gains are observed relative to learning-based baselines. Compared to LambdaMART, the policy improves NDCG@10 for 95.8% of cell lines, with a median increase of + 0.386. A similarly high proportion of improvements (95.8%) is observed relative to the regression-to-ranking baseline, with a median increase of + 0.312. These results indicate that directly optimizing a decision-oriented policy yields substantial advantages over surrogate ranking objectives, even when those objectives are learned from the same underlying data.

Importantly, improvements are observed across a wide range of baseline performance levels, including cases where baseline NDCG@10 is already high. This suggests that the policy not only corrects poor baseline rankings but also consistently refines drug prioritization across diverse cellular contexts.

These results demonstrate that the proposed policy-based framework provides robust and consistent improvements over both heuristic and learning-to-rank baselines in clinically relevant top-*k* settings.

## 6 Conclusion

In this work, we presented NetPolicy-RL, an offline reinforcement learning framework for pharmacogenomic drug prioritization that integrates network diffusion-based biological modeling with policy optimization. By directly optimizing ranking quality rather than point-wise drug response prediction, the proposed approach consistently improves decision-oriented performance across cancer cell lines.

Empirically, the learned policy achieves substantial reductions in per-cell regret, indicating that near-optimal drugs are more reliably identified among the top recommendations. Across multiple ranking metrics, including NDCG, regret, and oracle-based analyses, NetPolicy-RL consistently outperforms global ranking heuristics and learning-to-rank baselines. Ablation experiments further demonstrate that neither empirical response statistics nor network-derived features alone are sufficient; instead, their integration yields the strongest and most robust performance. These findings confirm that incorporating mechanistic network context meaningfully enhances drug prioritization beyond what can be achieved using response data alone.

The offline actor-critic optimization exhibits stable convergence and generalization across held-out cell lines, despite operating on a fixed dataset. This highlights the practicality of policy-based learning in pharmacogenomic settings, where online interaction and exploration are infeasible. Importantly, all biological quantities are computed deterministically prior to learning, preserving interpretability while enabling effective optimization.

Future work will focus on extending the network impact representation to learnable encoders capable of incorporating uncertainty-aware offline reinforcement learning objectives to further improve robustness. Overall, this study demonstrates that decision-centric, biologically informed policy learning provides a powerful and effective paradigm for drug prioritization in precision oncology.

